# The significance of mitochondrial DNA half-life to the lifespan of post-mitotic cells

**DOI:** 10.1101/2020.02.15.950410

**Authors:** Alan G Holt, Adrian M Davies

## Abstract

The proliferation of mitochondrial DNA (mtDNA) with deletion mutations has been linked to aging, and age related neurodegenerative conditions. In this study we model effect of mtDNA half-life on mtDNA competition and selection.

Individual cells effectively form a closed ecosystem containing a large population of independently replicating mtDNA. We would expect competition and selection to occur between wild type mtDNA and various mutant variants. There is a symbiotic relationship between the cell and the mitochondria, and unrestricted mtDNA replication would be detrimental to the host cell. Deletion mutations of mtDNA are relatively common and give a replication advantage to the shorter sequence, as this could be lethal to the host cell, we would expect to see differences in mtDNA replication in short and long lived cells.

In this paper, we use a computer simulation of mtDNA replication, where mtDNA sequences may undergo deletion errors and give rise to mutant species that can compete with the wild type. This study focuses on longer lived cells where the wild type mtDNA is expected to be more susceptible to displacement by mutants. Our simulations confirm that deletion mutations have a replication advantage over the wild type due to decreased replication time. Wild type survival times diminished with increased mutation probabilities. The relationship between survival time and mutation rate was non-linear; a ten-fold increase in mutation probability resulted in a halving in wild type survival time.

In contrast a modest increase in the mtDNA half-life had a profound affect on the wild type survival time in the presence of deletion mutants, thereby, mitigating the replicative advantage of shorter sequence mutations. Given the relevance of mitochondrial dysfunction to various neurodegenerative conditions, we propose that therapies to increase mtDNA half-life could be a therapeutic strategy.

## 1 Introduction

Research on mitochondrial DNA (mtDNA) has progressed rapidly since the seminal publications associating mtDNA mutation with human disease [24, 52]. Mutations in mtDNA are now known to be a causal factor in many pathologies [8, 30, 51].

Over the course of evolution, the majority of the genes present in the ancestral mtDNA have been transferred to the nuclear DNA [2, 9]. The resulting mtDNA is highly compact [3], as such, any deletions in the genome will disrupt gene function. Mammalian mtDNA encodes for 13 proteins of the electron transport chain, 22 tRNAs, and 2 rRNAs. The structure of the mitochondrial genome displays a remarkable economy, pocessing overlapping genes but no introns [48].

Given the close proximity of mtDNA to the electron transport chain, it will be exposed to damaging free radical species [53] and, compared to nuclear DNA, mtDNA has a somewhat limited capacity for repair [4]. This forms the basis of the mitochondrial free radical theory of aging [15, 22], which has been a dominant theory for several decades [21]. The free radical theory of aging, however, is not without critics [46]. It has been observed that the pattern of mutations seen in mtDNA is not what would be expected for oxidative damage [28] and may be more indicative of replicative error. Nevertheless, oxidative stress and free radicals are still widely regarded as a major source of damage to mtDNA. This damage has been implicated in aging and neurodegenerative disease [17, 45].

In contrast to nuclear DNA, individual cells possess between a few hundred to many thousands of copies of mtDNA. Also, unlike nuclear DNA which does not replicate during the lifetime of the cell, there is a relatively rapid turnover of mtDNA with a half-life dependent on the cell type. This half-life can vary from a few days to several weeks [18, 32, 36, 43]. Most of the research into the replication of mtDNA has been carried out in rodents and there is a paucity of data concerning the cell and tissue specific half-life of human mtDNA.

Data from various studies, predominantly in the rodent [6, 11, 39], are suggestive of a correlation between the life span of the cell type and the half-life of the mtDNA. For example, in the rodent, the mtDNA half-life is 20-30 days for long lived, post-mitotic cells such as neurons, but 8-11 days in short lived, fast replicating cells, such as hepatocytes or epithelial cells [18, 27, 31, 38, 43].

There are numerous studies that associate mitochondrial (dys)function with aging and age related pathologies [10, 26, 47]. It has been demonstrated that mitochondrial deletions (mtDNA_*del*_) increase in frequency with age [29]. Experiments indicate that these mtDNA_*del*_ may be instrumental in the process of aging [49], at least for humans; deletion mutations are rare in rodents [5].

The biological mechanisms that determine aging have been the subject of intense study for many decades, and remain unresolved, controversial, but well reviewed [13]. The proliferation of mtDNA_*del*_ is one factor in the process of aging and the pathology of neurodegenerative disease [19, 23, 42], but it is amenable to simulation and theoretical investigation.

Kowald and Kirkwood have demonstrated that mtDNA_*del*_ have a replicative advantage over wild type (mtDNA_*wild*_) [34, 35]. It is a reasonable supposition that, in post-mitotic somatic cells, there is an opportunity for selfish replicons [14], such as mtDNA_*del*_, to proliferate at the expense of the mtDNA_*wild*_ species [25, 50].

Blackstone and Kirkwood [1] predict that mtDNA_*wild*_ would be at a disadvantage in proliferating cells compared to mtDNA_*del*_ due to the need to expand the mtDNA population. However, high levels of deletion mutations are, generally, only found in post-mitotic cells [41]. We agree that mtDNA_*del*_ would have a strong selective advantage in proliferating cells but we would expect it be detected less often and have a low impact compared to postmitotic cells. In proliferating cells, rapid growth of the mtDNA_*del*_ population would lead to cell death and replacement. Unfortunately, post-mitotic cells are seldom replaced.

In this paper, we describe a computer simulation model of mtDNA cloning with deletion mutations. Initially we used it to reproduce the results of Kowald and Kirkwood [34, 35] that shows that deletions have a replicative advantage over wild types. The model was then used to explore the effects of half-lfe on the time taken for mtDNA_*del*_ to become dominant and drive the mtDNA_*wild*_ to extinction.

Current research suggests a link between mtDNA_*del*_ proliferation and the onset of neurodegenerative disease. Consequently, mtDNA half-life may not be long enough to support current expected human lifespan. Our results show that the proliferation of mtDNA_*del*_ is accelerated by its replicative advantage over mtDNA_*wild*_. However, a modest increase in half-life could have a significant effect on wild type survival and that therapies to increase mtDNA half-life, could be beneficial.

This research:

- Uses computer simulation to study mtDNA wild type survival in the presence mutations deletions.
- Confirms that deletion mutations have a replicative advantage over wild type due to shorter replication times, in agreement with Kowald and Kirkwood [34, 35] and validates our simulation model.
- Demonstrates that an increase in mtDNA half-life can mitigate the replicative advantage of mtDNA_*del*_ and extend mtDNA_*wild*_ times.

## 2 Simulator

In this section we describe the mitochondrial simulator. The simulator was written in Python [7], and relies heavily on object orientated programming methods. We create an organelle (software) object which comprise a population of mtDNA objects. Like their biological counterparts, the mtDNA objects can replicate and expire over time.

The simulation operates over a set period in discrete time intervals, *t*. The state of the system is updated in each time interval according a set of events that may occur in *t*. The simulation run is initialised at *t* = 0 with a population of mtDNA_*wild*_. In each iteration, new mtDNAs may be added to the population through cloning. MtDNAs are removed from the organelle through ageing and eventual expiry; thus, simulating the process of accumulation of damage and degradation. New species of mtDNAs result from random deletion mutations (mtDNA_*del*_). A mtDNA_*del*_ sequence is a fragment of the mtDNA_*wild*_ sequence. MtDNAs compete for space in an organelle with limited capacity. MtDNAs are prevented from cloning if the population exceeds capacity. Cloning is only reactivated when the population drops below capacity due to mtDNA expiry. Cloning is also controlled in response to ATP production (by mtDNA_*wild*_). Given a cell has certain energy requirements, an energy deficit will respond by activating cloning as a means of increasing ATP production. Conversely, cloning is deactivated when there is a surplus of energy.

### 2.1 Simulator Outline

The simulator undergoes an initialisation process by creating an organelle (software) object *O* which is populated with an initial set of mtDNAs (also software objects). The initial organelle state is denoted, thus:

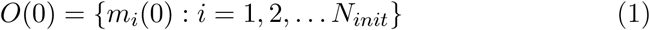

where *N*_*init*_ is the initial population size. *O*_*count*_(*t*) denotes the number of mtDNAs in the organelle at time *t. O*_*count*_(*t*) is incremented as new mtDNAs are cloned and decremented as they expire. We set the initial population of the organelle to *N*_*init*_ = 1, 000 mtDNA_*wild*_; thus, *O*_*count*_(0) = 1, 000. *C*_*max*_ is the capacity of the organelle such that *O*_*count*_(*t*) ≤ *C*_*max*_. For the purpose of this study, *C*_*max*_ = 2, 000.

Once the organelle and its initial population of mtDNAs has been created, the simulator loops through a number of time intervals *t* = 1, 2, *…, L*, where each iteration yields an organelle state *O*(*t*) and *L* is the run length of the simulation.

The time interval *t* represents 15 minutes of (real) time. During the life time of the organelle, mtDNAs will be both added and removed to form a new organelle state, *O*(*t* + 1). A brief outline of the steps performed by the simulator are given below: (pseudocode for the algorirthm is shown in Appendix (B)):

- Request ATP from the organelle’s pool. Record if there was (or was not) sufficient ATP in the pool to fulfil the request.
- For each mtDNA in the organelle, perform the following tasks:
  – If the TTL of the mtDNA is zero, then remove from *O*.
  – Update the state of the mtDNA:
    * Age the mtDNA (with probability *P*_*damage*_).
    * If the mtDNA is undergoing replication, decrement *busy*.
    * Produce ATP and add to ATP pool.
  – If the ATP request in deficit, then turn on cloning.
    * Clone (subject to conditions being met, see below).
    * Possibility of mutation.

Any mtDNA in *O*(*t*) with a *m.ttl* = 0 are deemed expired and not passed on to the next generation, *O*(*t* + 1).

Updating the state of an mtDNA *m* involves *ageing* it by decrementing its *time-to-live* (TTL), *m.ttl*. The TTL attribute is initialised, thus: *m.ttl* = *maxTTL* when *m* is created. Note, this is for newly cloned mtDNAs. When we create the organelle we populate it with mtDNAs and, for the initial population, we set *m.ttl* to a uniformly distributed random value between 1 and *maxTTL* so that mtDNAs appear to have different ages. In each iteration we run a Bernoulli trial to determine if the mtDNA sustains damage (or not). With a probability *P*_*damage*_, damage is inflicted on *m* by decrementing *m.ttl*. As a corollary, *m.ttl* remains unchanged with a probability 1 - *P*_*damage*_. The choice of *maxTTL* is somewhat arbitrary because we calibrate *P*_*damage*_ to yield a specified half-life for that *maxTTL*. In this paper we use a value *maxTTL* = 10. We compute *P*_*damage*_ using Markov model described in the appendix.

ATP is “produced” within the organelle for purpose of supplying energy to the cell in which it resides. ATP levels in the organelle are used as a signal to the mtDNA to increase or decrease their production of ATP. If there is a deficit in the ATP request, the ability of each mtDNA to clone is activated. Otherwise, it is deactivated. While a deficit in the ATP request will trigger the cloning function, it does not necessarily mean an mtDNA *m* will undergo replication. The model of ATP consumption/production is described in more detail in subsection 2.2. The cloning of *m* is summarized below:

- The success of a Bernoulli trial with probability *P*_*clone*_, Derivation of *P*_*clone*_ is described in the subsection below.
- *m.ttl >* 0: that is, the mtDNA needs to be alive.
- *m.busy* = 0: that is, the mtDNA is not currently undergoing cloning. If it is, *m.busy* ≤ 0.
- *O*_*count*_(*t*) ≤ *C*_*max*_, that is, the organelle capacity has *not* been exceeded.

In other studies, cloning probability is a function of the population size. According to Kowald *et al.* [34] it is assumed “that the replication probability declines with the total amount of existing mtDNAs” and that “some form of negative feedback exists that limits the production of new mtDNAs if sufficient exist”. Whereas, Kowald et al. [34] use a Verhulst logistic map to dynamically adjust the cloning probability according to population levels, we maintain a constant cloning probability, *P*_*clone*_ = 0.01. In our simulation, mtDNA populations are controlled by the ATP production (by wild types) and the maximum capacity of the organelle (*C*_*max*_). We do not; therefore, need to rely on continually adjusting a probability variable in relation to population size.

If all criteria for cloning are met, then a *child* clone of the original (parent) mtDNA results. The child; however, may mutate, in which case, the DNA sequence of the child is different to that of the parent. Mutation takes place upon the success of a Bernoulli trial of probability *P*_*mutate*_. If mutation occurs, the child’s DNA sequence is generated from a random fragment of its parent. Consequently, a mutant “clone” constitutes a new species of mtDNA where its DNA sequence is shorter than that of the parent. Obviously, if mutation does not take place, the child is the same species as its parent and their DNA sequences are identical.

Note that, over time, it is possible for mutations of mutations to occur (though this can be controlled in our simulator). We used various values for mutation probability, *P*_*mutate*_ ranging from 10^−3^ to 10^−2^. A realistic value is around 10^−3^, whereas values closer to 10^−2^ are probably, artificially high. Nevertheless, we were interested in the dynamics of the system as mutation probabilities increase.

Replication of an mtDNA takes time. For this reason, mtDNAs are placed in a *busy* state whilst undergoing replication. Prior to the start of the replication process, *m.busy* = 0, denoting an *idle* state. Upon entering the replication process, the busy attribute is set to *m.replTime*, where *m.replTime* is an attribute of *m* and denotes its replication time. The replication time of an mtDNA *m* is proportional to its size. The size of a mtDNA_*wild*_ sequences (*wildSize*) is 16,569 nucleotides. If *m* is a wild type, *m.size* = *wildSize* but if *m* is a mutant, then *m.size* is a fraction of *wildSize*. We compute an mtDNA_*del*_ replication time, thus:

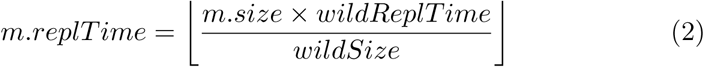

Associated with each mtDNA, *m*, is a DNA sequence, *m.seq*. MtDNAs with the same *m.seq* are of the same species. For the mtDNA_*wild*_ sequence, we use the NC_012920.1 Homo sapiens mitochondrion (this choice was arbitrary). For any mutant species, *m.seq* is a random fragment of NC_012920.1. While mitochondrial DNA sequences are small compared the nuclear DNA sequences, they still require several kilobytes of storage. As such, we do not want to store *m.seq* for every *m* that is created. For this reason, we use cryptography hashing functions to keep track of species of *m*. The attribute *m.type* is computed by applying a SHA224 hash algorithm to *m.seq*:

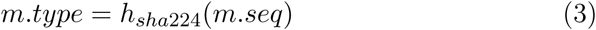

For each *m* we store the *m.type* (which is only 224 bytes) instead of *m.seq*. A unique list of *m.seq* values is stored in a table indexed by *m.type*. A lookup on the table is performed when *m.seq* is required. Using cryptography hashes for DNA sequences yields a considerable performance increase in terms of the simulator’s execution speed.

### 2.2 ATP Production and Consumption

The mitochondria are the organelles that are responsible for ATP production within the cell. The cell’s demand for energy in the form of ATP is modelled as an external “consumer” (to the organelle) that periodically requests a quantity of ATP. Each mtDNA_*wild*_ “produces” some proportion of the ATP which is added to a common ATP pool for the organelle. The consumer extracts ATP from the pool in order to fulfil the cell’s energy requirements, see Fig 1.

**Figure 1:**
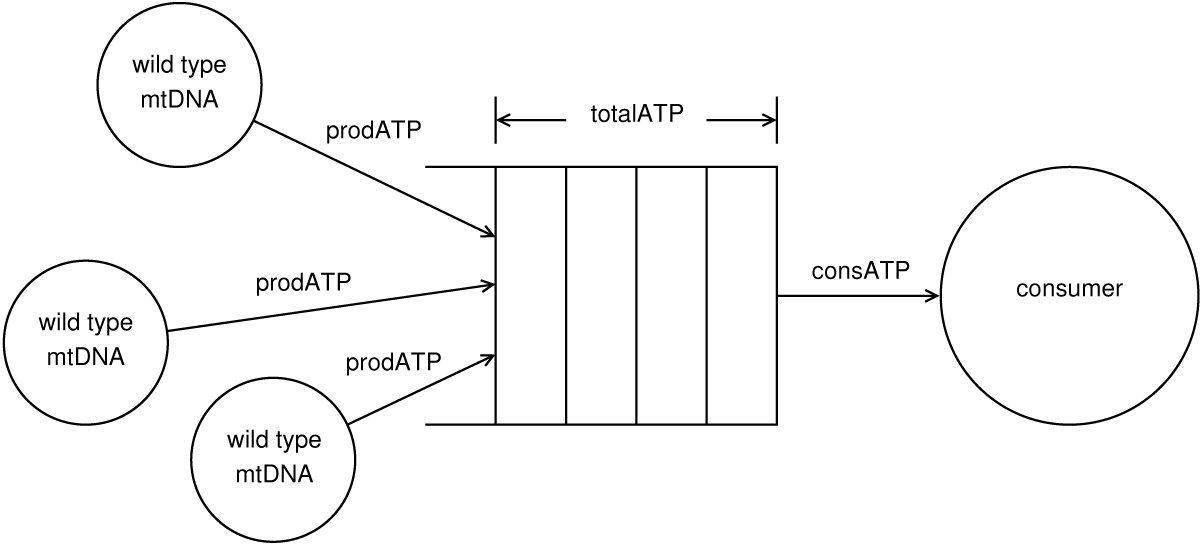
ATP production and consumption model.

In reality, we know that mtDNA sequences do not produce ATP directly and there is no common ATP pool. Nor does the external consumer exist as an actual biological entity. Our model is merely a convenient abstraction of the energy usage process within the cell. We use the ATP surplus/deficit to control behaviour within the organelle. If the ATP request by the external consumer is fulfilled, the ability to clone is turned off. If there are insufficient levels of ATP within the pool to fulfil the request, the ability to clone is turned on.

The attributes, prodATP and consATP are the amount of ATP that the *mtDNA*_*wild*_ produces and the amount consumed by the external consumer in each time interval, respectively.

## 3 Analysis and Results

In this section, we use our simulation to carry out two investigations, namely:

- The replicative advantage of deletion mutations have over wild type mtDNA. Kowald et al. [33–35] have already demonstrated that deletion mutations have a replicative advantage. The purpose of this investigation is, primarily, validate over simulation and verify it is giving expected results.
- The effects of half-life on wild type survival times.

### 3.1 Replicative Advantage of Deletion Mutations

We consider the survival times of the mtDNA_*wild*_ species in the presence of mtDNA_*del*_ in a single mitochondrial organelle, where mtDNA_*del*_ maybe multiple species. The mtDNA replication times vary according to the size of the mtDNA sequence. For mtDNA_*wild*_ the replication time is two hours. As mtDNA_*del*_ sequences are a fraction of the size of mtDNA_*wild*_, their replication times are proportionately less. In order to investigate if shorter replication times give mtDNA_*del*_ an advantage over mtDNA_*wild*_, we ran a control experiment where replication times were kept constant regardless of size of the mtDNA; that is, mtDNA_*wild*_ and mtDNA_*del*_ take the same time to replicate. We refer to these two replication strategies as *variable* and *constant* and we compare the survival time of the mtDNA_*wild*_ between the two.

We ran the simulation 30 times for each value of *P*_*mutate*_ = 1 × 10^−3^, 2 × 10^−3^, *…*, 1 × 10^−2^ and recorded the time that elapsed until the mtDNA_*wild*_ became extinct (a summary of the simulation parameters are given in Table (1)). However, given that the simulation run length was limited to 30 years, it was possible that the mtDNA_*wild*_ species survived until the end of simulation. In this situation, the survival time for mtDNA_*wild*_ was recorded as 30 years and marked as a *censored* data point.

**Table 1:** Simulation parameters

| Parameter | Value | Comment |
| --- | --- | --- |
| $T$ | 30 years | Simulation run length |
| $P_{damage}$ | 0.0101 | Equiv. 10 day half-life |
| $maxTTL$ | 10 | |
| $C_{max}$ | 2000 | Capacity of organelle |
| $N_{init}$ | 1000 | Initial population |
| $wildReplTime$ | 2 hours | Wild type replication time |

Given the possibility of censored data points, we adopt survival analysis methods [16] to analyse our results. We fit Kaplan-Meier survival curves which give the probability function for the survival time of the mtDNA_*wild*_ species. Curves were computed for each value of *P*_*mutate*_ and replication strategy, namely, variable and constant. The Kaplan-Meier survival curves are shown in Fig 2. For the control simulations (constant replication times), where there is no replicative advantage for mtDNA_*del*_, wild type survival times are governed by random drift. In contrast, where there is a replicative advantage, there is a detrimental effect on wild type survival.

**Figure 2:**
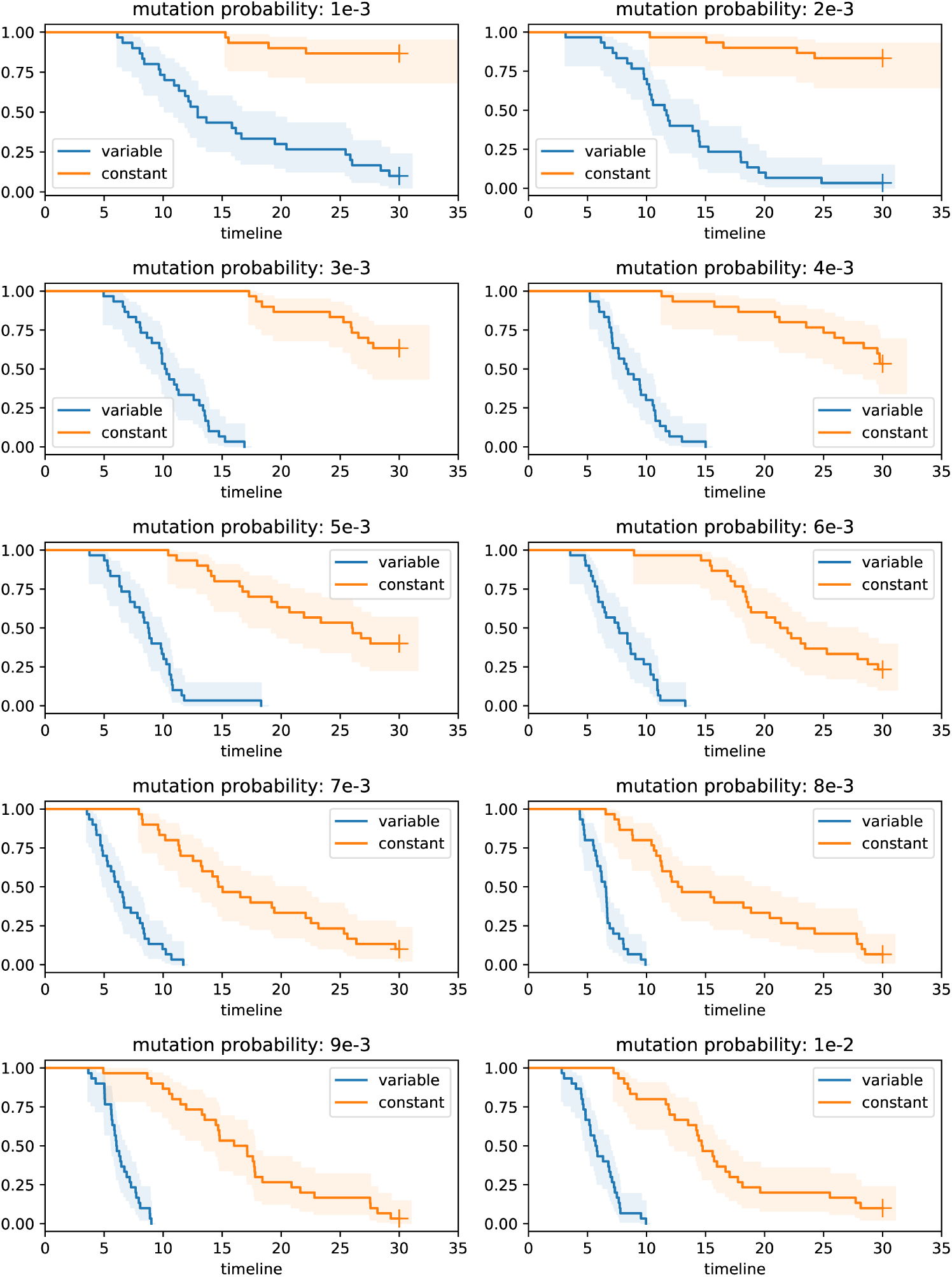
Survival Analysis (timeline in years)

The graphs show that the likelihood of mtDNA_*wild*_ survival is lower where replication times vary according to mtDNA sequence size compared to constant replication times (for all mutation probabilities). Note that, the + symbol at the end of a curves, denotes censored data points.

Figure 3 shows the median^1^ survival time versus mutation probability. We fit the following non-linear model:

**Figure 3:**
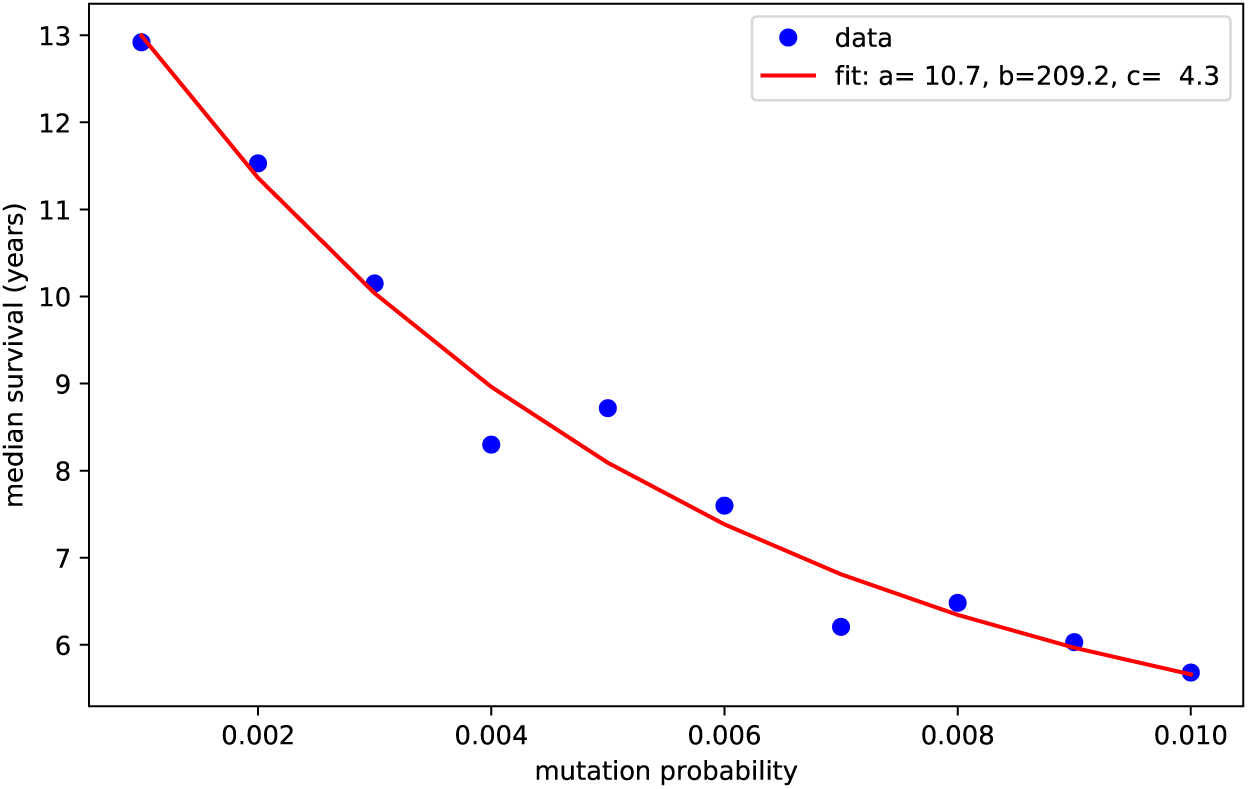
Median mtDNA_*wild*_ survival times.

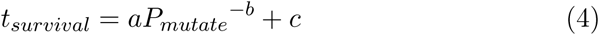

where *t*_*survival*_ is the median survival time of the mtDNA_*wild*_ species (and *P*_*mutate*_ is the mutation probability). The model parameter values are *a* = 10.7, *b* = 225.8 and *c* = 4.8. This result confirms expected behaviour, however, what is notable about our result, is that, for a ten fold increase in mutation probability, the survival time is merely halved.

The *constant* replication time strategy was only performed as an experimental control; while survival times do seem to correlate with mutation probability, these results are; somewhat, artificial.

### 3.2 Effects of Half-life on Wild Type Survival

The primary purpose of this study was to investigate the effect of half-life on mtDNA_*wild*_ survival time. We ran the simulator as before, but with mtDNA half-lives of 10, 15, 20, 25 and 30 days. We increased the run length of the simulation to 70 years (approximate average life span of a human host). Just as with the previous experiment, simulations were run 30 times for each half-life level and we used a mutation probability of *P*_*mutate*_ = 10^−3^. The simulation parameters are summarised in Table (2).

**Table 2:**
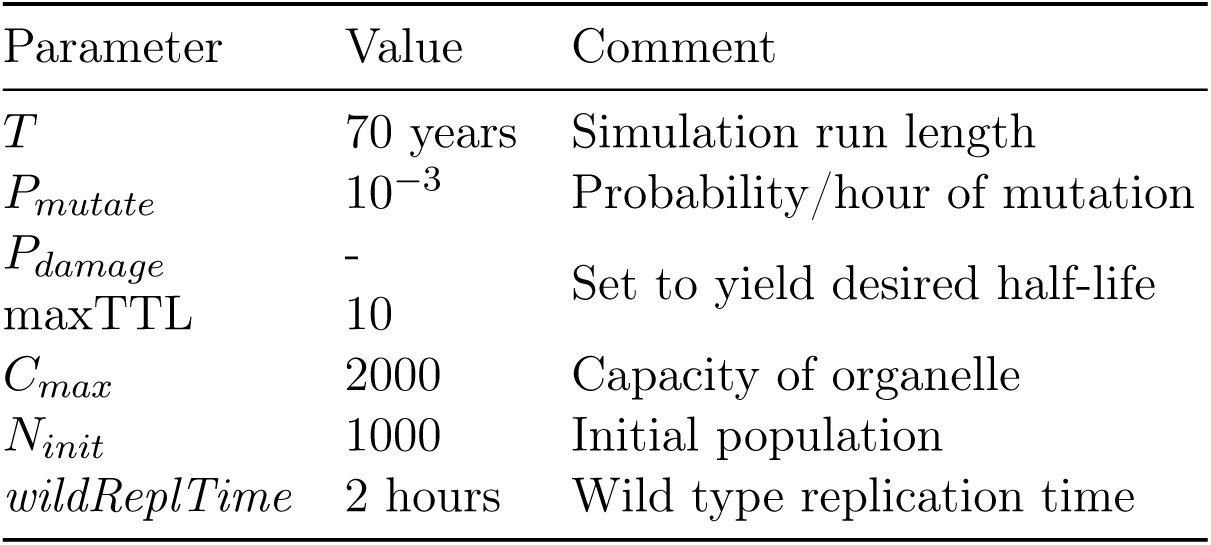
Simulation parameters for half-life experiment

The median mtDNA half-life times are shown in Table 3 and the survival curves are shown in Fig 4. It is apparent that there is a marked increase in mtDNA_*wild*_ survival time for quite modest increases in half-life. For a 30 day half-life, the Kaplan-Meier function yielded a median value of ∞; that is, it was unable to derive a finite median value given the number of censored data points. Nevertheless, we can see that the likelihood of the mtDNA_*wild*_ species surviving well beyond 70 years, is quite high. Figure 5 shows the mtDNA_*wild*_ median survival time for half-lives, for which there was a finite value.

**Table 3:** mtDNA_*wild*_ survival time versus half-life

| Half life<br>(days) | Median survival<br>time (years) |
| --- | --- |
| 10 | 14.5 |
| 15 | 23.8 |
| 20 | 52.4 |
| 25 | 64.6 |
| 30 | ∞ |

**Figure 4:**
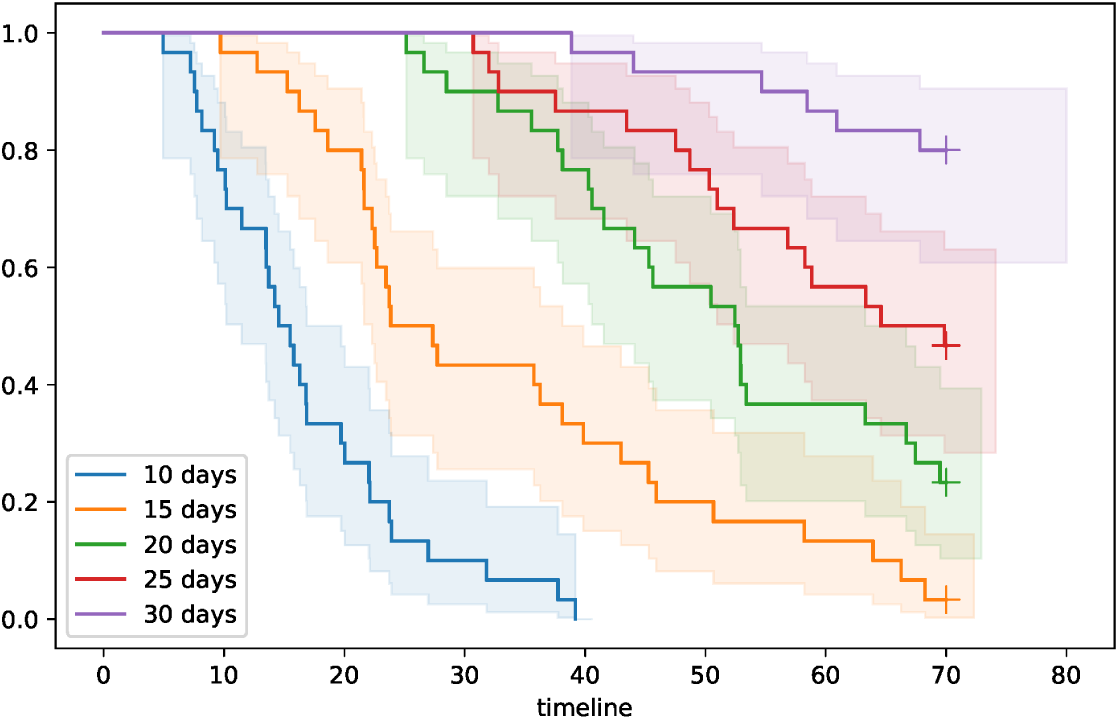
Survival curves for mtDNA half-lives 10, 15, 20, 25 and 30 days (timeline in years)

**Figure 5:**
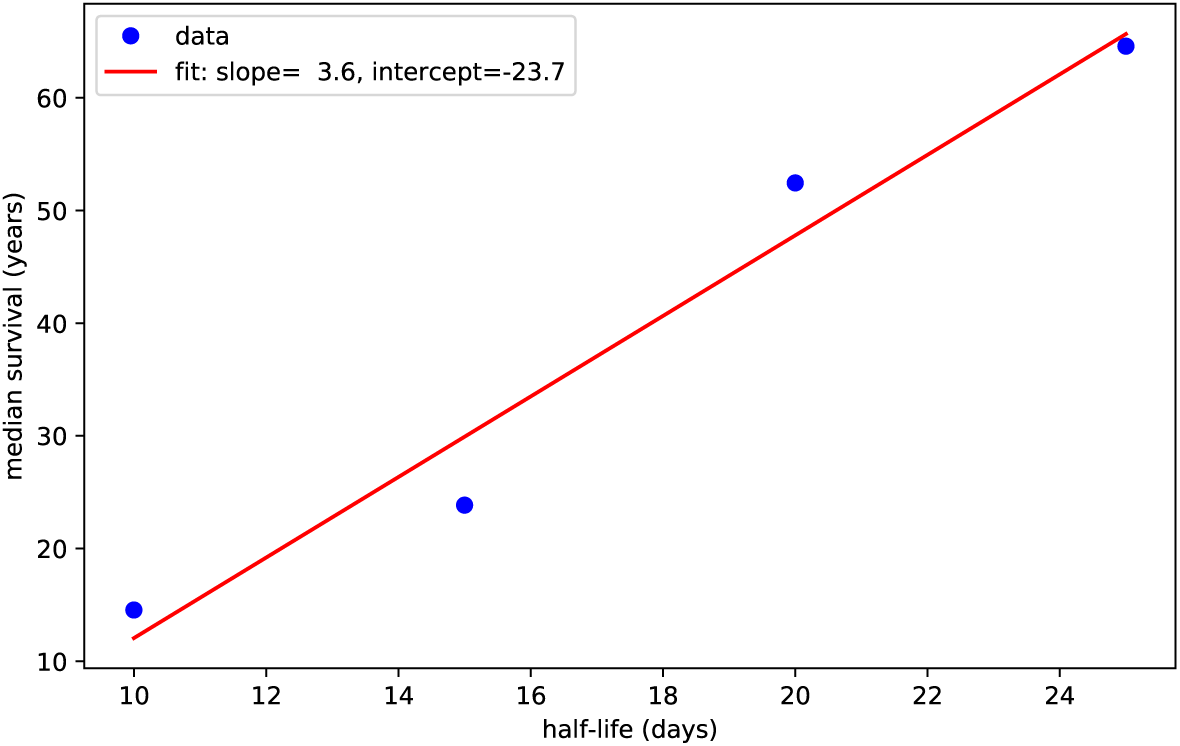
Survival time versus half-life.

We fit a linear regression curve to this data with slope 3.6 and intercept −23.7. From this model, we can predict that the survival time for the missing 30 day half-life is 84.6 years. While current half-life levels in post-mitotic cells may have been sufficient for most of our evolutionary history, they are not long enough to support current expected human lifespans. Consequently, there is a significant probability that, people living into their 80s and beyond are at a high risk of neurodegenerative pathologies. Therapies that target extending mtDNA half-life could have a profound effect on reducing this risk. Furthermore, this increase could be quite modest; according to our model, a half-life of 35 days would yield a median survival time of 102.3 years.

## 4 Discussion

The mitochondrion within a given cell can be considered as a single “virtual” organelle [40, 44]; essentially a closed ecosystem containing competing replicons. Within this system there is the potential for mtDNA_*del*_ to proliferate at the expense of the mtDNA_*wild*_ species [34], and drive the mtDNA_*wild*_ to extinction within the lifespan of the cell. This process will be of greater consequence for organs, such as the brain which is composed of post-mitotic cells with little capacity for regeneration.

While the biological process involved in the proliferation of mtDNA_*del*_ is undoubtedly complex, the problem is amenable to computer modeling and simulation. To investigate the impact of mtDNA half-life we developed a computer simulation of mitochondrial cloning and mutation. Our results demonstrate that, the half-life of the mitochondrial genome has a profound effect on the population dynamics of mtDNA, and on the competition between mtDNA_*wild*_ and mtDNA_*del*_ species. A commonly cited value for the half-life of mitochondrial DNA is 10 days [34]; however, empirical observations suggest that the half-life can be much longer, especially in post-mitotic cells [43].

Incorporation of tritiated thymidine into the mtDNA has been used in combination with autoradiography to measure the half-life of mtDNA in various cell types in the mouse, it is clear that the mtDNA half-life is not uniform across tissues [18, 31]. While the actual values for mtDNA half-life may not directly translate from mouse to human, it seems plausible that the half-life of human mtDNA also varies between cell types. The half-life of mtDNA in relatively short lived cells, such as epithelial cells or hepatocytes, is 8-12 days [31], whereas, in post-mitotic cells, such as neurons, the half-life of the mtDNA is 20-30 days [18, 31]. The results from our simulation show that mtDNA_*del*_ have a replicative advantage over mtDNA_*wild*_ due to their smaller size.

The graphs in Figs 6 and 7 show the mtDNA populations for two particular simulation runs, namely, *variable* replication time (replication times proportional to mtDNA size) and *constant* replication time (control), respectively. The blue line in the graphs (referenced in the legend) show the wild type population, while the other colours represent a species of mutant. It can be seen that, while a number of mutant species populate the organelle, one particular mutant species dominates. These examples demonstrate clonal expansion within a mitochondrion. Clonal expansion, as exhibited in these examples, was common in our simulation runs, however, it was not always the case. Sometimes, there was no outright single mutant species that dominated. Unfortunately, a thorough analysis of heteroplasmy is outside the scope of this paper, but is a subject of future research.

**Figure 6:**
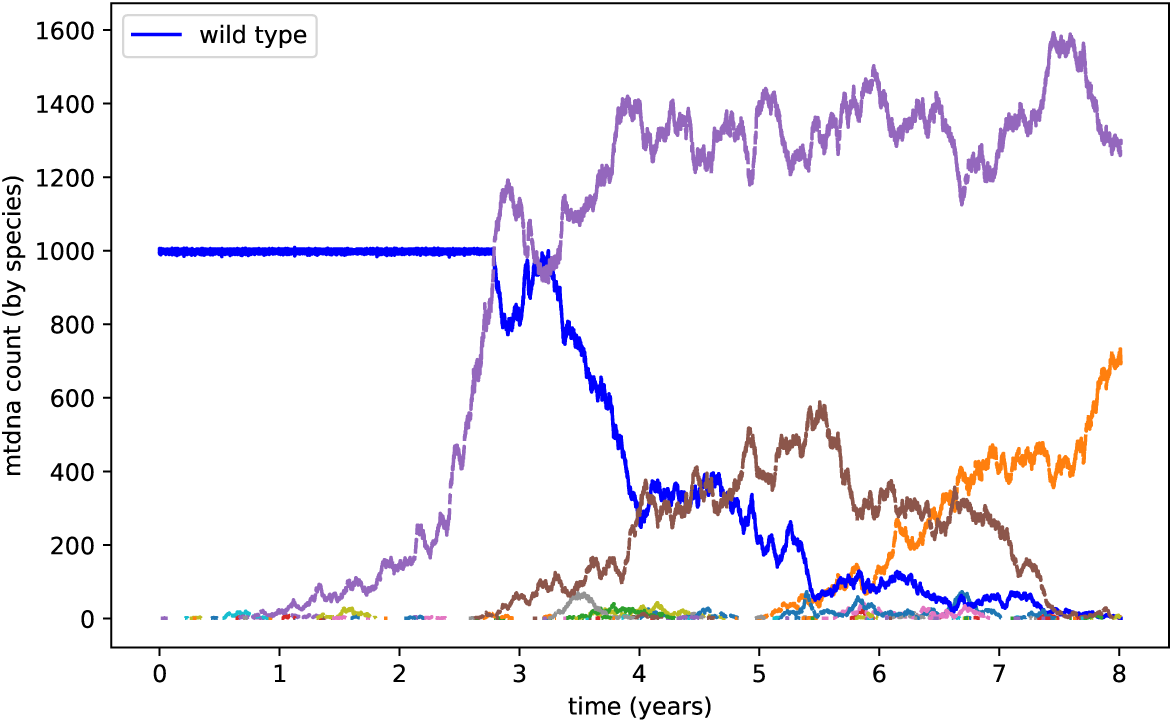
mtDNA species timeseries for *variable* replication (*P*_*mutate*_ = 10^−3^, half-life 10 days)

**Figure 7:**
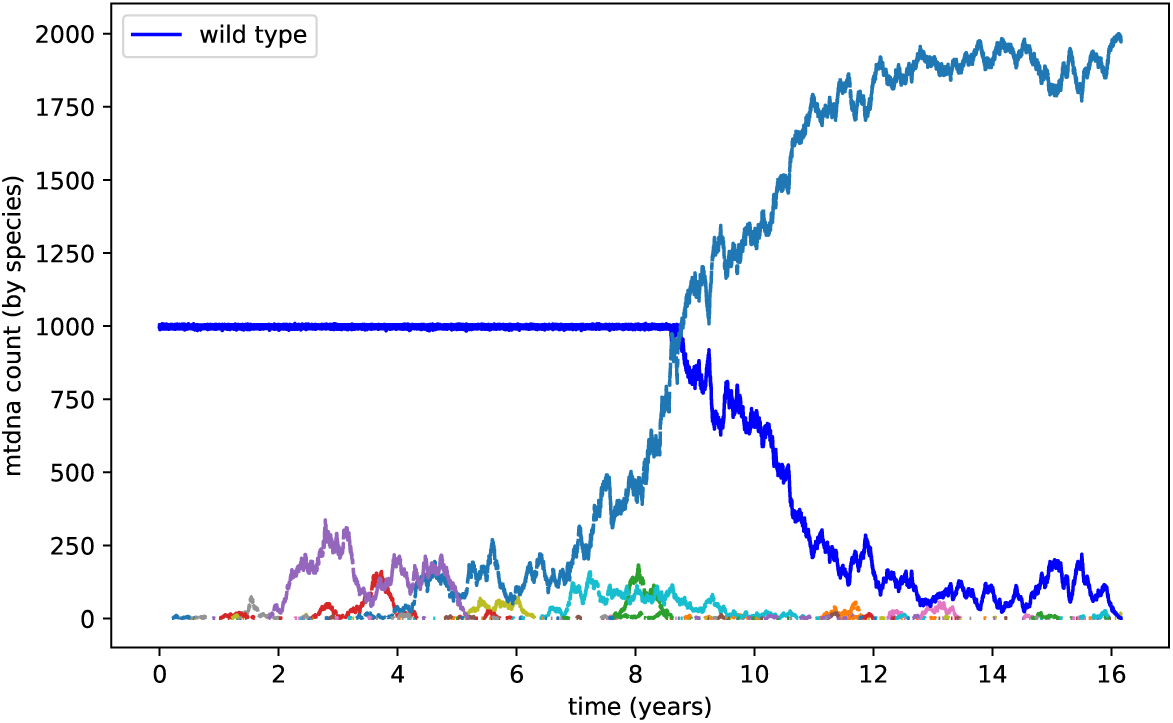
mtDNA species timeseries for *constant* replication (*P*_*mutate*_ = 10^−3^, half-life 10 days)

We ran simulations over a range of mutation probabilities and this result was consistent across the range; this result confirms common expectations. Also, as expected, mtDNA_*wild*_ survival times diminished with increasing mutation probability. Unexpectedely, these times diminished slowly; for a tenfold increase in mutation probability, there was only a halving of survival time.

If the predominant cause of mutation is free radical damage, it is unlikely that increasing the level of antioxidants would be an effective means of reducing the level of mtDNA_*del*_ [37]. Whereas, if the predominant cause is replication error [28] an increase in proofreading would; presumably, increase survival time. The fact that polymerase-*γ* does not have more accurate proofreading would suggest that; it is not necessary, not possible, or is energetically prohibitive. We propose that it is not necessary, as an increase in survival time can be achieved with an increase in the mtDNA half-life.

For a cell type with a short lifespan of weeks or months, such as skin fibroblasts or blood cells, rapid proliferation of mtDNA_*del*_ is of little consequence as the host cell would have died and been replaced before any mutants became dominant. For cells, such as neurons that are post-mitotic, and need to last a lifetime, proliferation of mtDNA_*del*_ will be catastrophic.

For post-mitotic cells, such as neurons, that are not normally replaced or have a low rate of turnover; loss of functional mitochondria, during their expected working life, would have serious consequences. The mtDNA halflife in neuronal cells in the mouse is greater than 20 days [31]. If we assume that the mtDNA in human neuronal cells has an equally long half-life, it is expected that it would take more than 70 years for mtDNA_*del*_ to become dominant.

Modern medicine has led to significant changes in human demographics. Between 1933 and 2014 the mean lifespan for women increased from 63 to 81 [12], unfortunately the greatest rate of decrease in white and grey matter in the brain occurs between the ages of 65 and 80 [20].

Based on these results we posit that therapies that extend the mtDNA half-life could slow mtDNA_*del*_ proliferation, thereby, extending mtDNA_*wild*_ survival.

## 5 Conclusions

We simulate a cell that regulates cloning of mtDNA according to ATP production by mtDNA_*wild*_. The cell needs to maintain a level of mtDNA_*wild*_ for its energy needs. This can become problematic in the presence of mtDNA_*del*_. As the population of mtDNA_*del*_ (selfishly) grows, the organelle will eventually reach its capacity; thus, inhibiting all mtDNAs’ ability to clone (irrespective of species). In the event of an ATP deficit, the cell is prevented from increasing ATP production by means of cloning. mtDNAs can only resume cloning once the population drops below the capacity limit due to mtDNAs expiring. Once cloning has been reactivated, mtDNA_*wild*_ and mtDNA_*del*_ then compete for the available capacity. As we have seen, the shorter replication times of mtDNA_*del*_; due to their smaller sequence sizes, give them an advantage in this competition. Proliferation of mtDNA_*del*_ eventually brings about the extinction of mtDNA_*wild*_. These results are in agreement with previous publications.

In this paper we investigated the effect of mtDNA half-life on mtDNA_*wild*_ survival times. Our results show that higher half-lives slowed mtDNA_*del*_ proliferation, thus, increasing a cell’s lifespan. An implication of this study is that, the half-life of mtDNA is optimized for, and proportional to, the half-life of the host cell. For example, in the cells of the central nervous system; which are post-mitotic and have minimal potential for replacement or renewal. In these cells displacement of mtDNA_*wild*_ by mtDNA_*del*_ will lead to cell death and progressive loss of function.

Advances in medicine, nutrition, and other social factors mean that, a lifespan of greater than eighty years is not unusual. Increased longevity, however, is not without costs; the risk of neurodegenerative diseases increases with age. At eighty years dementia of some form will affect one in five people; we propose that the relatively long half-life of mtDNA in post-mitotic cells, such as neurons, may function so as to delay the proliferation of mtDNA_*del*_, but is probably optimal for a maximal lifespan shorter than we currently enjoy. Our results show that a 35 day half-life extends mtDNA_*wild*_ survival time beyond 100 years. Thus, even a modest increase in mitochondrial DNA half-life has the potential to delay or even prevent, mitochondrial or neurodegenerative pathologies. Therapies that could delay the onset of dementia, even by a few years, could have a significant impact.

## Acknowledgements

The authors would like to thank Dr Chi-Yu Huang and Dr Daniel Ives for their valuable feedback.

## Appendix A Markov Model for Half-life

In this appendix, we present a Markov model to derive a value of *P*_*damage*_ that yields a given half-life. The model comprises a series of states where each state represents the *m*_*ttl*_ = *n* value of the mtDNA. The Markov model transitions from state *n* to state *n* - 1 with a probability of *p* = *P*_*damage*_. Consequently, the mtDNA remains in the same state (that is, *m*_*ttl*_ is unaffected) with probability 1 - *p*, The transition probability matrix **P** for *maxTTL* = 10 is given by:

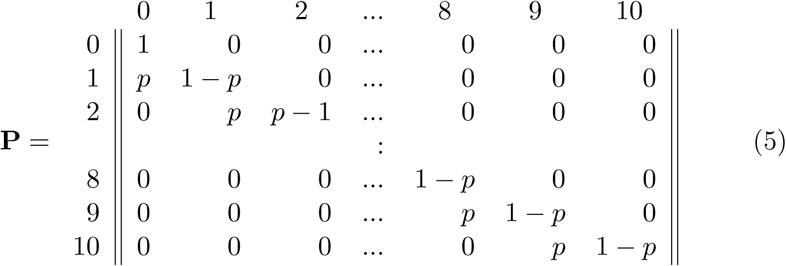

The initial state *π*^0^ is:

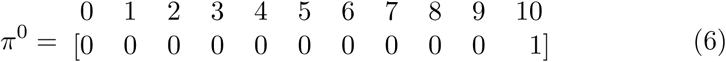

We demonstrate the Markov model by computing the mutation probability for a half life of 10 days. Given that the iteration interval is 15 minutes, then 10 days is 960 intervals (24 × 4 × 10). We ran the Markov model for various values of *p* until we achieved result close to:

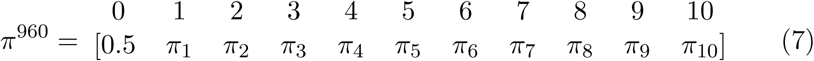

When *π*[0]^960^ ≈ 0.5, approximately half of the mtDNAs have expired as they have a TTL value *m*_*ttl*_ = 0. Similarly, approximately half of the population is still alive, that is, *m*_*ttl*_ *>* 0: We found *p* = 0.0101 yielded a half life of 10 days.

## Appendix B Simulation Pseudocode

This appendix contains the pseudocode for the simulator. Table 4 describes the data structures and functions used in the pseudocode.

**Table 4:**
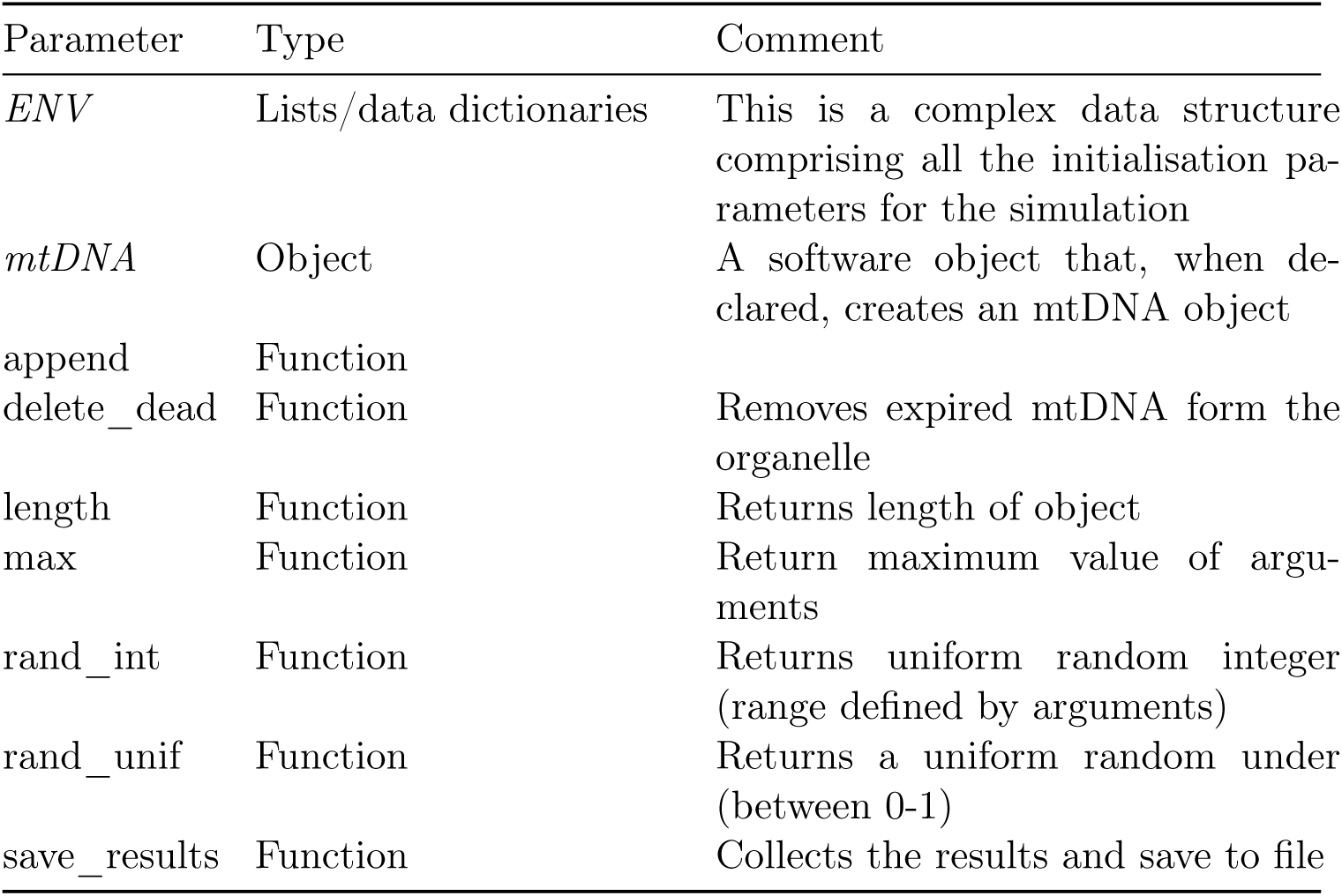
Simulation data structures and Functions

### Algorithm 1 Run Simulator

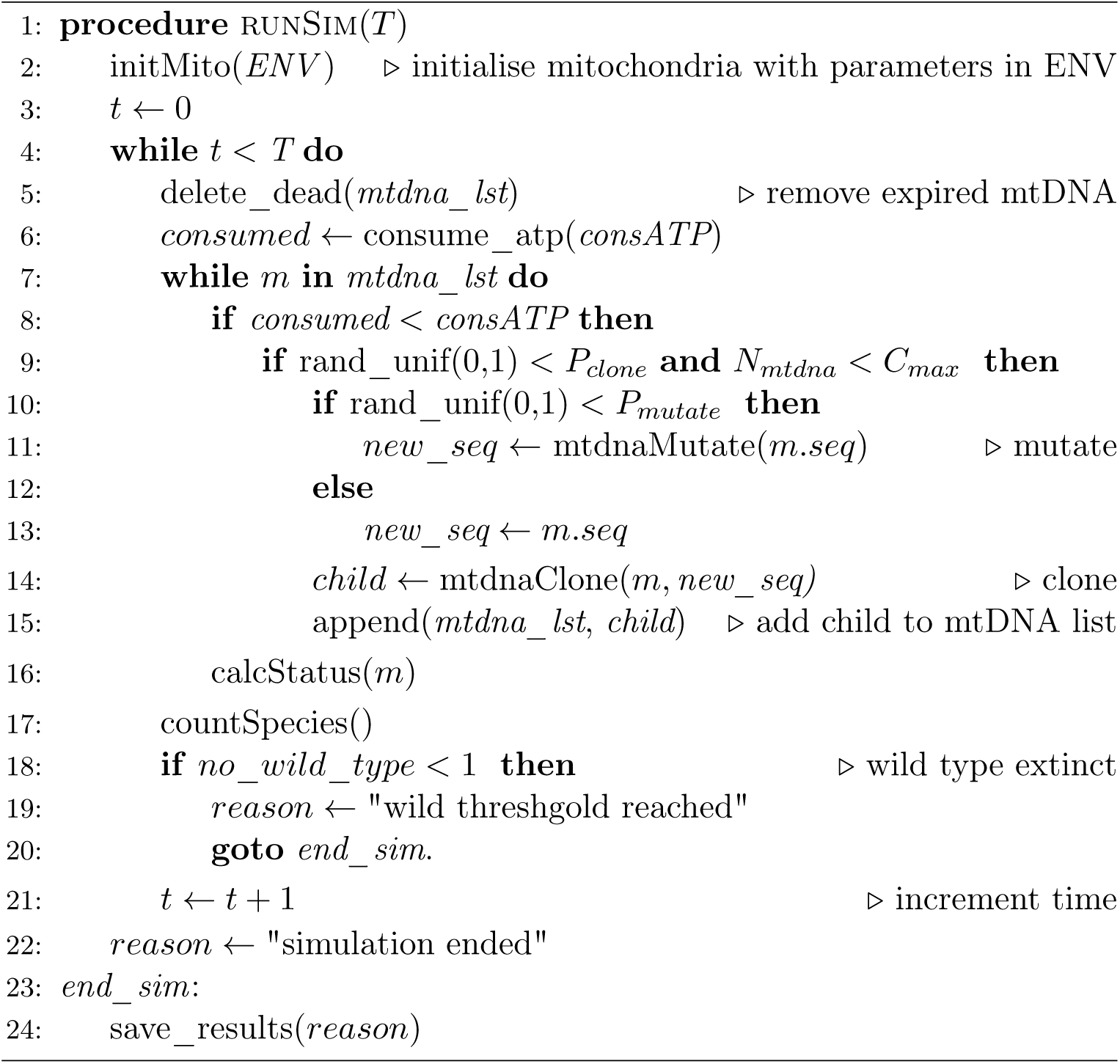

### Algorithm 2 Initialise Mitochondrial Organelle

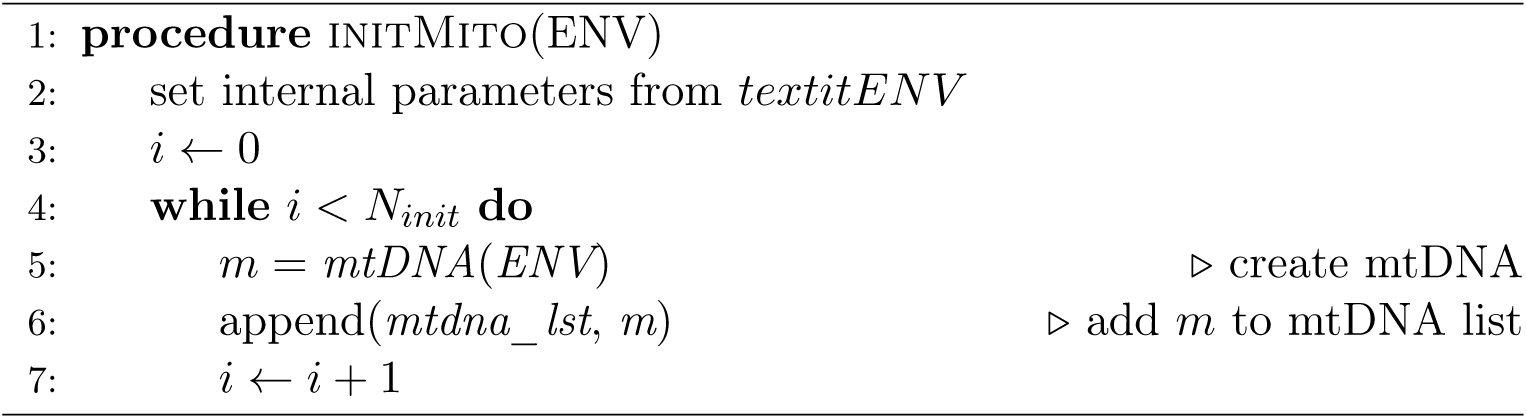

### Algorithm 3 Calculate mtDNA status

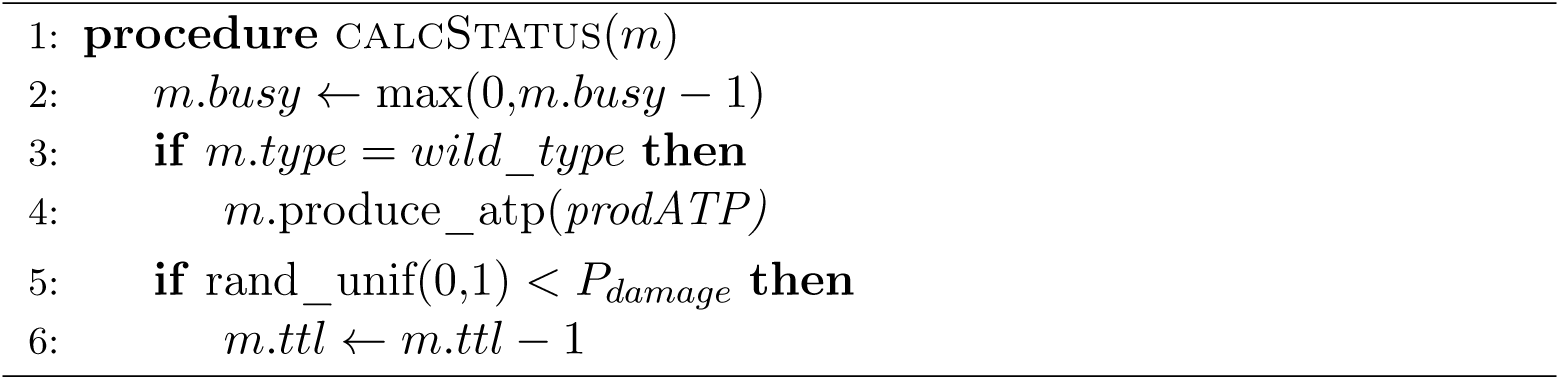

### Algorithm 4 Clone mtDNA

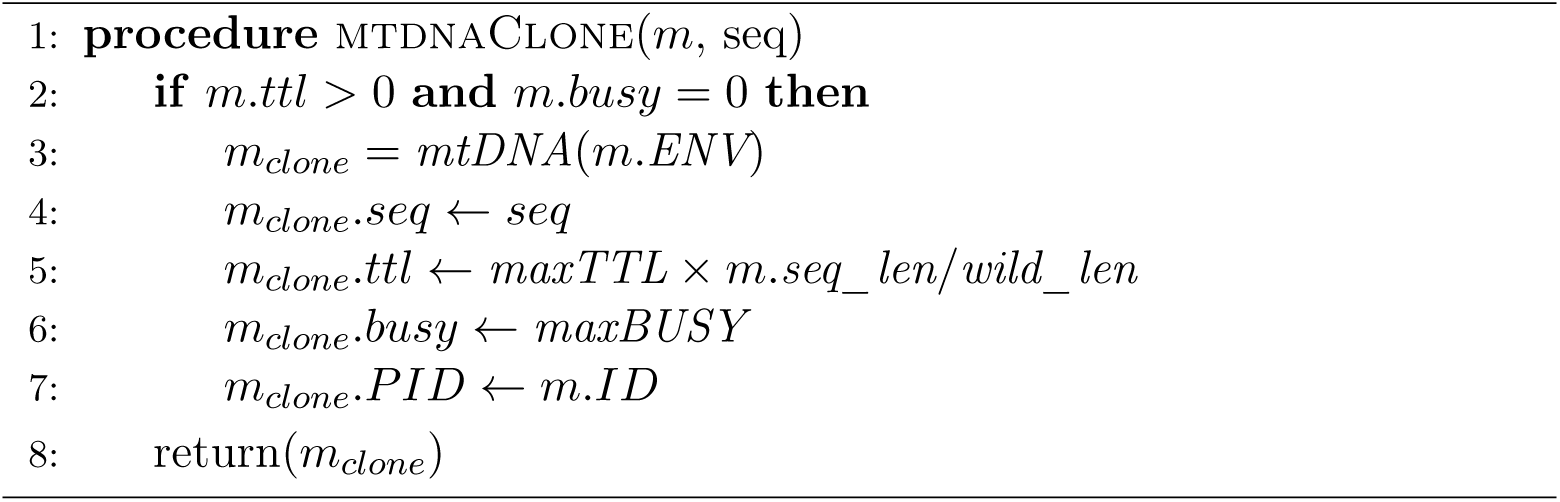

### Algorithm 5 Mutate mtDNA

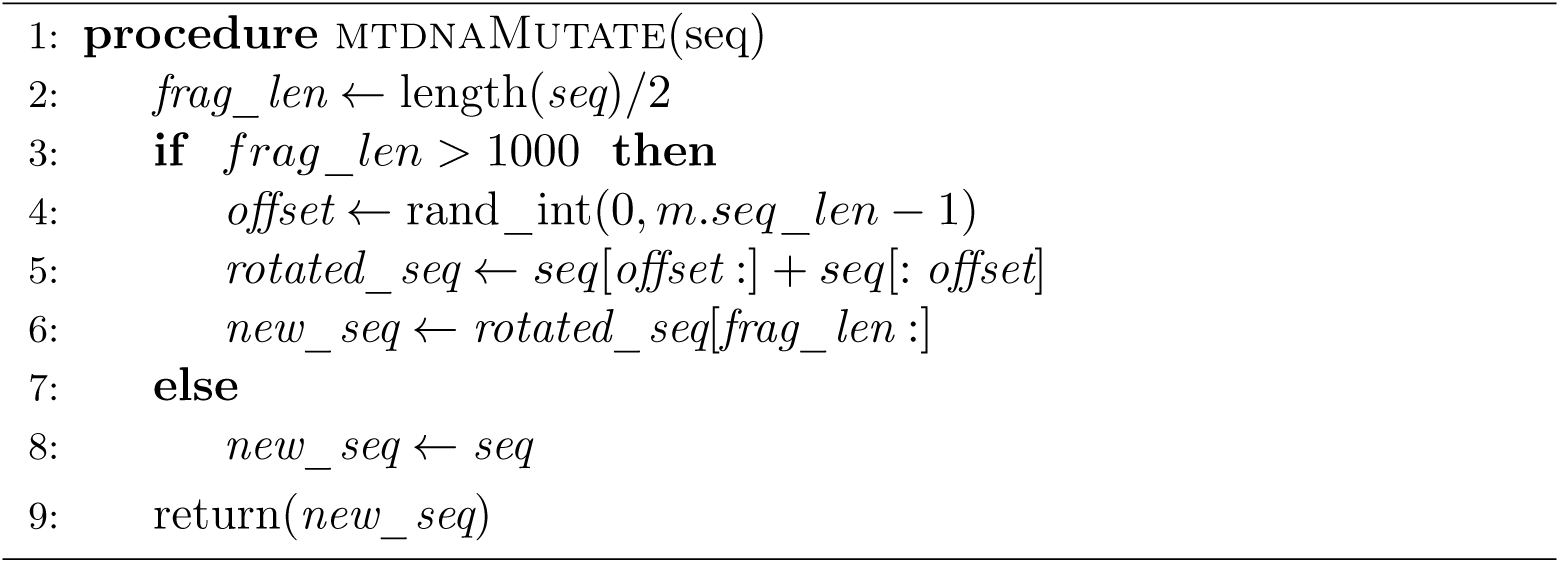

## Footnotes

1 The Kaplan-Meier it function also yields a median value

